# A Noninvasive Molecular Clock for Fetal Development Predicts Gestational Age and Preterm Delivery

**DOI:** 10.1101/212910

**Authors:** Thuy T. M. Ngo, Mira N. Moufarrej, Marie-Louise H. Rasmussen, Joan Camunas-Soler, Wenying Pan, Jennifer Okamoto, Norma F. Neff, Keli Liu, Ronald J. Wong, Katheryne Downes, Robert Tibshirani, Gary M. Shaw, Line Skotte, David K. Stevenson, Joseph R. Biggio, Michal A. Elovitz, Mads Melbye, Stephen R. Quake

**Affiliations:** Departments of Bioengineering and Applied Physics, Stanford University and Chan Zuckerberg Biohub, Stanford CA USA; Department of Epidemiology Research, Statens Serum Institute, Copenhagen, Denmark; Department of Statistics, Stanford University, Stanford CA USA; Department of Pediatrics, Stanford University School of Medicine, Stanford CA USA; Center for Women’s Reproductive Health, Department of Obstetrics and Gynecology, University of Alabama at Birmingham, Birmingham, AL USA; Maternal and Child Health Research Center, Department of Obstetrics and Gynecology, University of Pennsylvania School of Medicine, Philadelphia, PA USA; Department of Medicine, Stanford University School of Medicine, Stanford CA USA; Department of Biomedical Data Sciences, Stanford University School of Medicine, Stanford CA USA

## Abstract

We performed a high time-resolution, longitudinal study of normal pregnancy development by measuring cell-free RNA (cfRNA) in blood from women during each week of pregnancy. Analysis of tissue-specific transcripts in these samples enabled us to follow fetal and placental development with high resolution and sensitivity, and also to detect gene-specific responses of the maternal immune system to pregnancy. We established a “clock” for normal pregnancy development and enabled a direct molecular approach to determine expected delivery dates with comparable accuracy to ultrasound, creating the basis for a portable, inexpensive fetal dating method. We also identified a related gene set that accurately discriminated women at risk for spontaneous preterm delivery up to two months in advance of labor, forming the basis of a potential screening test for risk of preterm delivery.

## Introduction

Understanding the timing and programming of pregnancy has been a topic of interest for thousands of years. In antiquity, the ancient Greeks had surprisingly detailed knowledge of various details of stages of fetal development, and developed mathematical theories to account for the timing of important landmarks during development, including delivery of the baby (*1–3*). In the modern era, biologists have amassed detailed cellular and molecular portraits of both fetal and placental development. However, these results relate to pregnancy in general, and have not directly led to molecular tests that might enable monitoring of fetal development and predicting delivery for a given pregnancy. The most widely-used molecular metrics of development are the determination of the levels of human chorionic gonadotropin (HCG) and alpha-fetoprotein (AFP), which can be used to detect conception and fetal complications, respectively; however, neither molecule (either individually or in conjunction) has been found to precisely establish gestational age (GA) (*4, 5*).

Owing to the lack of a useful molecular test, most clinicians use either ultrasound imaging or a woman’s estimate of her last menstruation period (LMP) to establish GA and an approximate delivery date. However, these methods are imprecise and inadequate for gauging preterm birth, which is a substantial source of morbidity and mortality. Moreover, inaccurate dating can misguide the assessment of fetal development even for normal term pregnancies, which has been shown to ultimately lead to unnecessary induction of labor and Cesarean-sections, extended postnatal care, and/or increased medical expenses (*6–9*).

Understanding the progression of a normal pregnancy would contextualize abnormal phenotypes like preterm birth and establish methods to monitor pregnancy for signs of abnormalities or risk of preterm birth. Approximately 15 million neonates are born prematurely every year worldwide (*10*). As a major annual cost for the United States upwards of $26.2 billion (*11*), premature birth is the leading cause of neonatal death and the second most common cause of childhood death under the age of 5 years (*12*). Complications continue later into life as preterm birth is a leading cause of life years lost to ill health, disabilities, or early deaths (*13*). Soberingly, two-thirds of preterm deliveries occur spontaneously, and the only predictors are a previous history of a preterm birth, multiple gestations, and/or vaginal bleeding (*11*). Efforts to find a genetic cause have had only limited success (*14–16*) Thus far, most efforts have relied on epidemiological studies and the identification of environmental risk factors (*17*).

Here, we report a high time-resolution, longitudinal study of normal pregnancy development through measurements of cell-free RNA (cfRNA) in blood from women during each week of pregnancy. By quantifying gene expression in placental, fetal, and maternal tissues through cfRNA measurements, we open a window into the phenotypic state of a pregnancy. We have previously shown that the use of tissue-specific genes enables direct measurements of tissue health and physiology, which are concordant with the known physiology of pregnancy and fetal development at low time-resolution (*18*). Now we establish a high-resolution “clock” for normal pregnancy development and enable a direct molecular approach to determine time to delivery and GA. We demonstrated that cfRNA samples from both the second (13 to 24 weeks) and third (25 to 40 weeks) trimesters of pregnancy can predict expected delivery dates with comparable accuracy to ultrasound, creating the basis for a portable, inexpensive dating method. While this dating approach was validated only in normal pregnancies, we were also able to identify and validate biomarkers that accurately discriminated women at risk of spontaneous preterm delivery up to two months in advance of labor, thus indicating which women were at risk for deviating from the normal programmed developmental clock and forming the basis of a potential screening test for risk of preterm delivery.

## Results and Discussion

For our initial study, we recruited 31 pregnant Danish women, each of whom agreed to donate blood on a weekly basis, resulting in a total of 521 plasma samples (Figure 1). All women delivered at term, defined as a GA at delivery of ≥ 37 weeks, and their medical records showed no unusual health changes during pregnancy (Table 1). Each sample was analyzed by highly multiplexed real-time PCR using a panel of genes that were chosen to be specific to the placenta, fetal organ-specific tissues, or the immune system (*18*).

**Figure 1:**
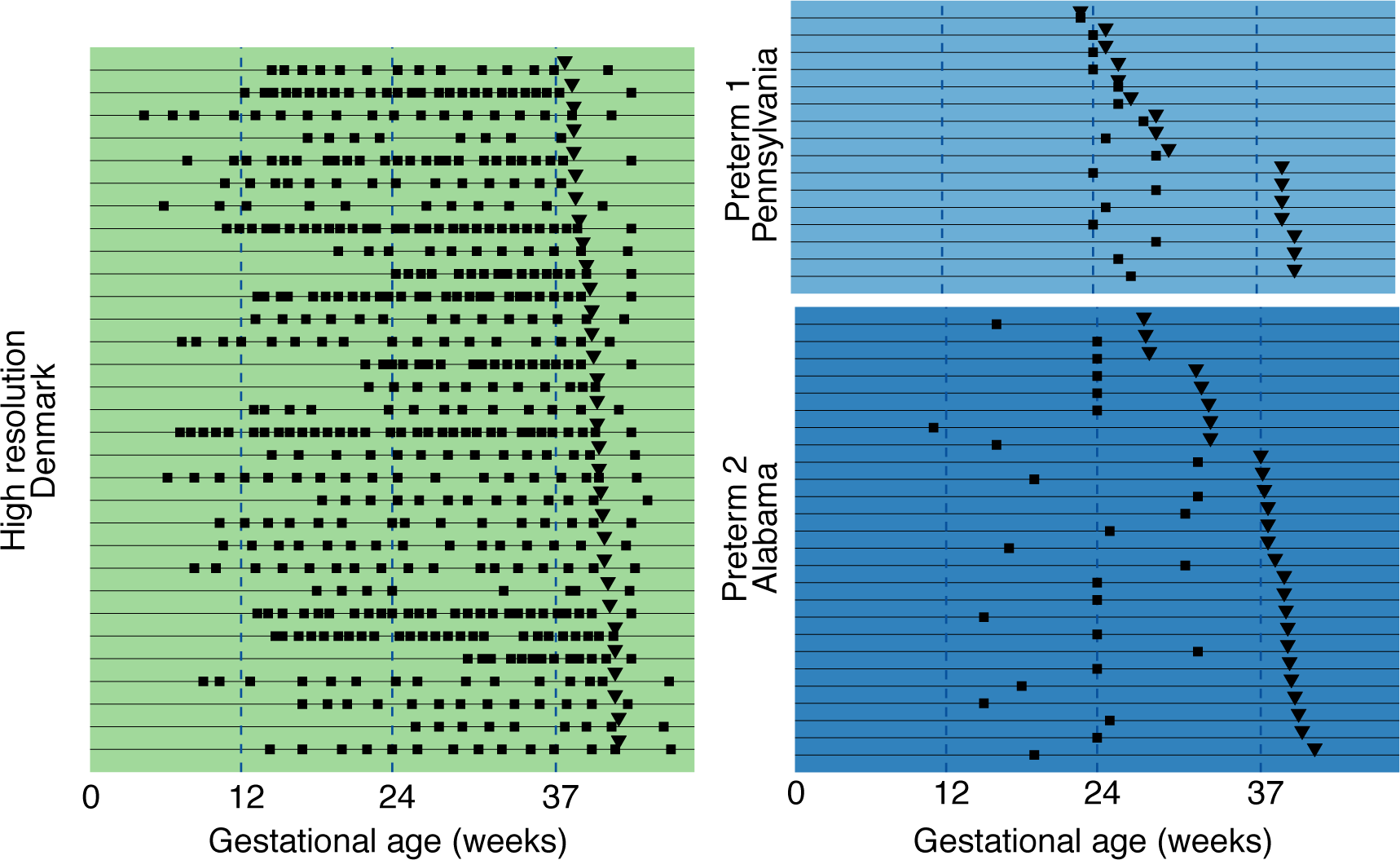
Sample collection timelines from Denmark, Pennsylvania, and Alabama. Squares, inverted triangles, and lines indicate sample collection, delivery dates, and individual women, respectively.

**Table 1:** Patient and pregnancy characteristics.

| **Demographics** | **Denmark**<br>**(n=31)** | **Pennsylvania (n = 16)** |  | **Alabama (n = 26)** |  |
| --- | --- | --- | --- | --- | --- |
|  | *At-term*<br>(n = 31) | *Preterm*<br>(n = 9) | *At-term*<br>(n = 7) | *Preterm*<br>(n = 8) | *At-term*<br>(n = 18) |
| Age (years, mean±SD) | 29.9±3.2 | 23.6±5.7 | 23.7±3.4 | 23.9±2.8 | 25.8±4.4 |
| Parity (% nulliparous) | 19 (61.3) | 4 (44.4) | 3 (42.9) | 0 (0) | 0 (0) |
| BMI (kg/m<sup>2</sup>, mean±SD) | 22.1±3.6 | 24.4±5.1 | 31.9±5.7 | 28.9±10.5 | 28.6±7.0 |
| Ethnicity (% Hispanic) | 0 (0) | 0 (0) | 0 (0) | 0 (0) | 0 (0) |
| Caucasian (%) | 31 (100) | 0 (0) | 0 (0) | 0 (0) | 1 (6) |
| African-American (%) | 0 (0) | 9 (100) | 7 (100) | 8 (100) | 17 (94) |
| Gestational age at delivery (weeks, mean±SD) | 40±1.2 | 26.7±2.3 | 39.4±0.5 | 30.8±2.5 | 38.7±1.2 |
| Mode of delivery |  |  |  |  |  |
| Spontaneous | 23 (74.2) | 5 (55.6) | 7 (100) | 7 (88) | 16 (29) |
| Cesarean-section | 8 (25.8) | 4 (44.4) | 0 (0) | 1 (12) | 2 (11) |
| Gender (% male) | 14 (45.2) | 4 (44.4) | 4 (57.1) | 5 (63) | 10 (56) |
| Birthweight (kg, mean±SD) | 3.6±0.6 | 1.0±0.3 | 3.4±0.4 | 1.7±0.7 | 3.1±0.4 |

The average time course of gene expression differed by gene function (Figures 2A, S1). Placental and fetal genes, defined as organ-specific developmental transcripts, showed a clear increase through the course of pregnancy. Some of these genes plateaued before delivery and one (CGB) decreased from a peak found in the first trimester. Both placental and fetal organ-specific cfRNAs were not present or barely detected after delivery, which supports their predominantly fetal-specific origins. Immune genes, which are dominated by the maternal immune system, but may also include a fetal contribution, had a more complex trajectory, but, in general, showed changes in time with measurable baselines early in pregnancy and after delivery. We then calculated the correlation between estimated gene transcript counts across all genes and all pregnancies (Figure 2B) and discovered that genes within each set (i.e. placental, immune, and fetal) were highly correlated with each other (Pearson correlation r = 0.66 (placental), 0.69 (immune), 0.62 (fetal)). Moreover, we found that placental and fetal genes also showed a moderate degree of cross correlation (r = 0.45), suggesting that placental cfRNA may provide an accurate estimate of fetal development and GA throughout pregnancy.

**Figure 2:**
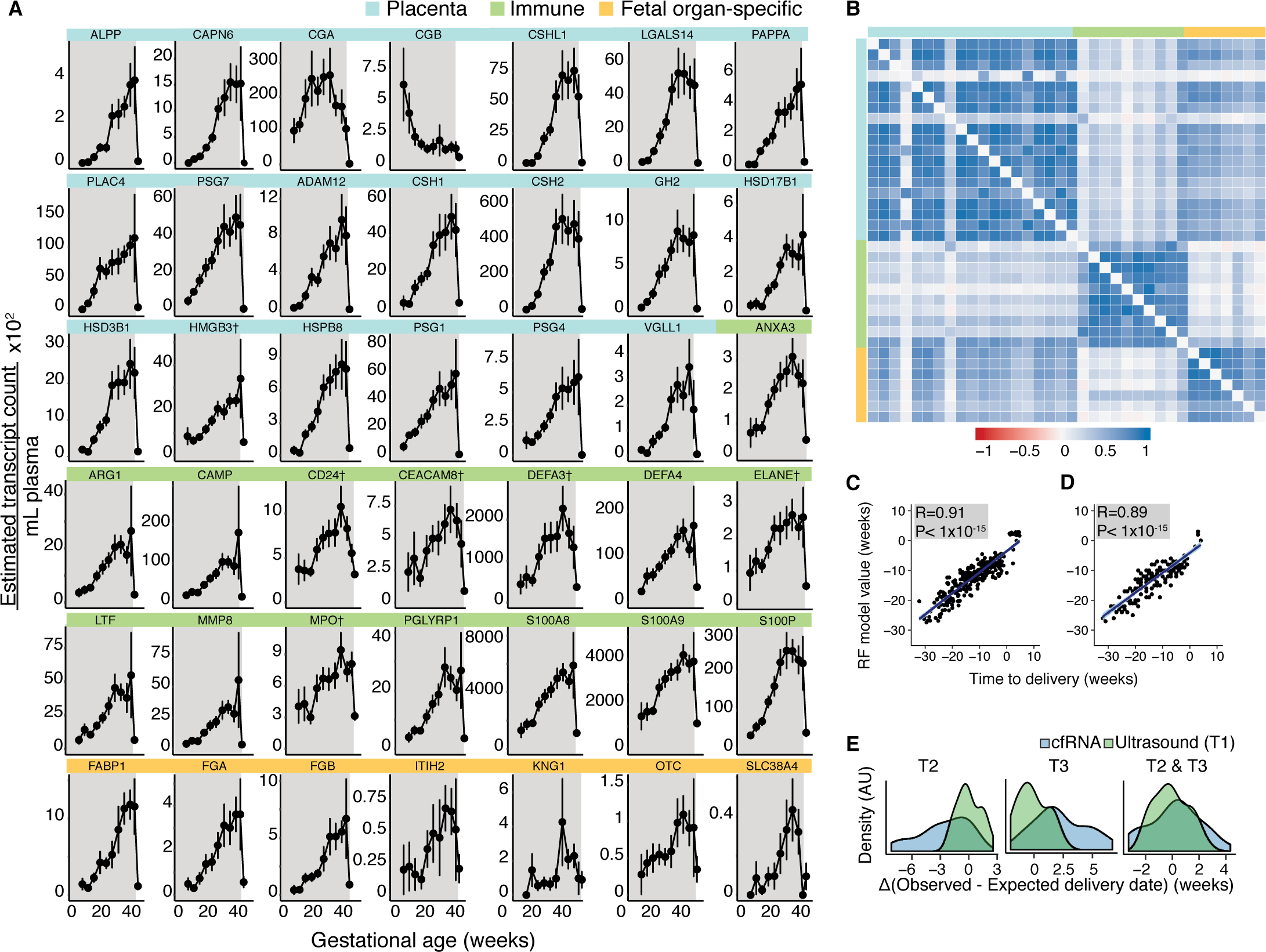
(A) Gene-specific inter-woman monthly means±standard error of the mean (SEM) plotted over the course of gestation (shaded in gray) for placental, immune, and fetal organ-specific genes. † represents genes for which data for only 21 women were available. (B) Correlation between gene-specific estimated transcript counts. Genes are listed in the same order as (A) omitting genes for which data was only available for 21 women. Color bar represents Pearson correlation r values. (C-D) Solid blue lines and shadings indicate linear fits and 95% confidence intervals, respectively. (C) Random forest model prediction of time to delivery for training data (n = 21, r = 0.91, P < 1 × 10^−15^, cross-validation). (D) Random forest model prediction of time to delivery for validation data (n = 10, r = 0.89, P < 1 × 10^−15^). (E) Comparison of expected delivery date prediction by ultrasound versus cfRNA.

Building upon these observations, we sought to build a more accurate predictor of GA using a machine learning model with cfRNA measurements as the primary features. We trained a random forest model on cfRNA data from 21 women across 24 time points (n = 306) and were able to show that a subset of 9 placental genes provided more predictive power than using the full panel of measured genes (Figure S2). Using these 9 genes (CGA, CAPN6, CGB, ALPP, CSHL1, PLAC4, PSG7, PAPPA, and LGALS14), we accurately predicted the time from sample collection until delivery (Pearson correlation r = 0.91, P < 1×10^−15^), which was an objective criterion independent of ultrasound-estimated GA (Figure 2C). Our model’s performance improved significantly over the course of gestation (root mean squared error (RMSE) = 6.0 (1^st^ trimester, T1), 3.9 (2^nd^ trimester, T2), 3.3 (3^rd^ trimester, T3), 3.7 (post-partum, PP) weeks). Remarkably, our model performed equally well (r = 0.89, P < 1×10^−15^) on a separate cohort of 10 Danish women (RMSE = 5.4 (T1), 4.2 (T2), 3.8 (T3), 2.7 (PP) weeks) (Figure 2D). We also built a separate model to predict GA (as estimated by ultrasound) and using the same 9 placental genes, the model performed comparably well both in training (r = 0.91, P < 1×10^−15^) and in validation (r = 0.90, P < 1×10^−15^) (Figure S3).

The random forest model selects placental genes as most predictive of time from sample collection until delivery and GA. Although, several of these genes showed similar changes in gene expression over time, their detection rate early in pregnancy varied. The redundant expression profiles of these genes may improve accuracy at early timepoints, when both placental and fetal cfRNAs were low and led to drop-out effects. As cfRNA increased during gestation, the accuracy of the model improved. This is in contrast to the efficacy of ultrasound dating, which relies on a constant fetal growth rate, an assumption that deteriorates over time (*7, 19*).

Further investigating the drivers of the model revealed several markers with known roles during pregnancy. CGA and CGB, the two most important features together with CAPN6, behaved differently from other genes in the model. CGA and CGB are the two subunits of HCG, known to play a major role in pregnancy initiation and progression and are involved in trophoblast differentiation (*20*). The trend observed for these two genes was consistent with what is known from HCG levels during pregnancy (*21*). Free CGB and PAPPA are also used as biochemical markers for risk of Down syndrome in the first trimester (*22*). Other genes selected by the model were related to trophoblast development (LGALS14, PAPPA).

We then used our model to estimate expected delivery dates from samples taken during the second, third, or both trimesters (Figure 2E). We found that 32% (T2), 23% (T3), 45% (T2 and T3), and 48% (T1 ultrasound) of all 31 Danish women delivered within 14 days of their expected delivery dates (Table 2). Similarly, prior studies reported that under normal circumstances, women delivered within 14 days of the expected date with 57.8% accuracy using ultrasound and 48.1% using LMP (*7*). Our results are not only comparable to ultrasound measurements, but also use a method that is inexpensive and more easily ported to resource-challenged settings.

**Table 2:**
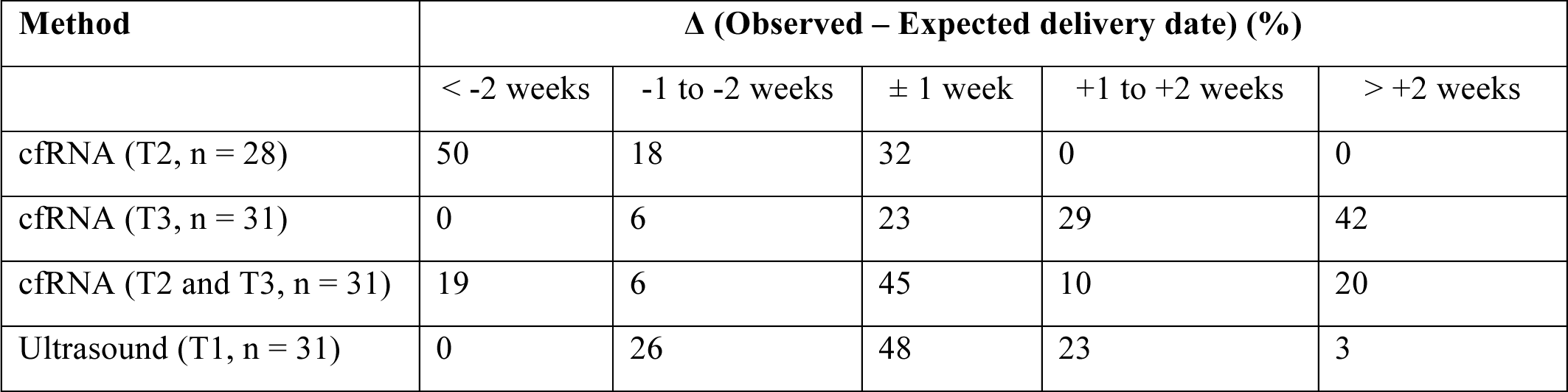
Comparing the distribution of expected delivery date predictions using cfRNA measurements from only the second (T2), third (T3), or both (T2 and T3) trimesters and ultrasound measurements from the first trimester (T1).

| Method | $\Delta$ (Observed – Expected delivery date) (%) | | | | |
| --- | --- | --- | --- | --- | --- |
| | < -2 weeks | -1 to -2 weeks | $\pm 1$ week | +1 to +2 weeks | > +2 weeks |
| cfRNA (T2, n = 28) | 50 | 18 | 32 | 0 | 0 |
| cfRNA (T3, n = 31) | 0 | 6 | 23 | 29 | 42 |
| cfRNA (T2 and T3, n = 31) | 19 | 6 | 45 | 10 | 20 |
| Ultrasound (T1, n = 31) | 0 | 26 | 48 | 23 | 3 |

While the first generation clock model was able to predict GA and time of delivery for normal pregnancies, we were also interested in testing its performance for predicting preterm delivery. We therefore used two separately-recruited cohorts, one collected by the University of Pennsylvania and the other by the University of Alabama at Birmingham, representing populations at elevated risk for spontaneous premature delivery to test model performance on an abnormal phenotype (Figure 1, Table 1). We discovered that while the model validated performance for full-term pregnancies (RMSE = 4.3 weeks) in these cohorts, it generally failed to predict time until delivery for preterm deliveries (RMSE = 10.5 weeks) (Figure S4). This suggests that the model’s content was reflective of a normal term pregnancy and may not account for the various outlier physiological events that may lead to preterm birth. In other words, from a molecular perspective, the premature fetus does not appear to have reached full gestation, and therefore preterm birth is likely not caused by fetal or placental “overmaturation” signals. This conclusion is supported by the observation that pharmacological agents designed to stop or slow down uterine contractions prevent only a small number of preterm deliveries (*23, 24*).

To further investigate this question and identify genes, which discriminate a spontaneous preterm from a full-term delivery, we performed RNA-sequencing (RNA-Seq) on plasma-derived cfRNA collected from women who delivered at full-term (n = 7) and preterm (n = 9) in a preterm-enriched cohort (Pennsylvania) (Figure 1, Table 1). Analysis of RNA-Seq data indicated that nearly 40 genes could separate term from preterm births with statistical significance (P < 0.001, See Methods) (Figure 3A). We then created a PCR panel with the highest scoring candidate preterm biomarkers and other immune and placental genes. We confirmed that the differential expression observed in our RNA-Seq analyses was also observed with this qPCR panel (Figure S5).

**Figure 3:**
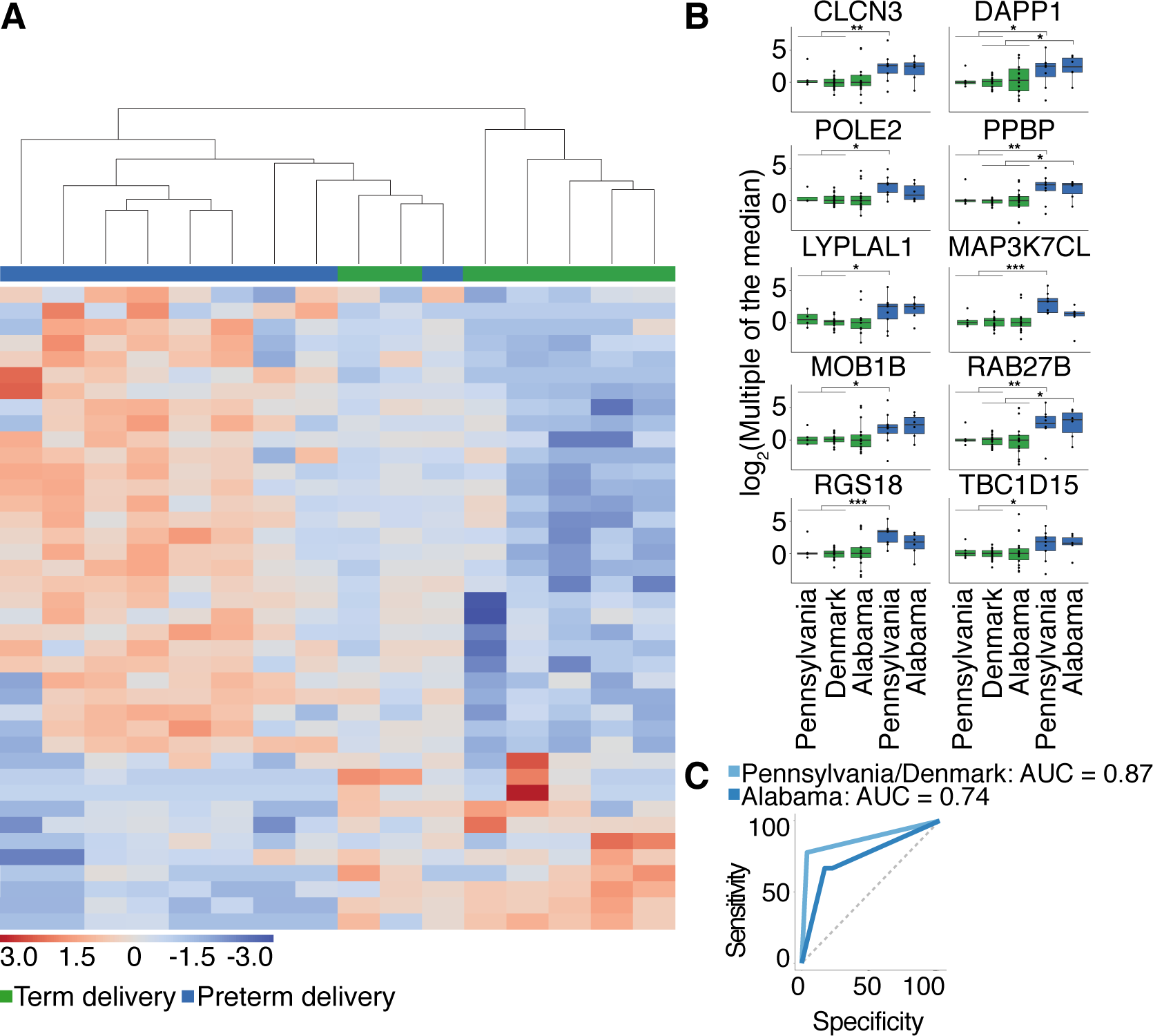
(A) RNA-Seq performed on samples from Pennsylvania highlights 40 differentially-expressed genes (P < 0.001) between preterm and term deliveries. Color bar represents z-scores. (B) Individual plots of 10 genes identified and validated in an independent cohort from Alabama, which accurately predicted preterm delivery in unique combinations of 3. All P-values reported are calculated using the Fisher exact test (FDR < 5%). *, **, and *** indicate significance levels below 0.05, 0.005, and 0.0005, respectively. (C) Predictive performance of 10 validated preterm markers in unique combinations of 3. Area under the curve (AUC) values are highlighted both for the discovery (Pennsylvania and Denmark) and validation (Alabama) cohorts.

When used in unique combinations of three (Table S1), the top ten genes from the panel (CLCN3, DAPP1, POLE2, PPBP, LYPLAL1, MAP3K7CL, MOB1B, RAB27B, RGS18, and TBC1D15) (FDR ≤ 5%, Hedge’s g ≥ 0.8) (Figure 3B), accurately classified 7 out of 9 preterm samples (78%) and misclassified only 1 of 26 full-term samples (4%) from both Pennsylvania and Denmark with a mean AUC of 0.87 (Figure 3C). These 10 genes in combination also showed successful validation in an independent preterm-enriched cohort from Alabama, accurately classifying 4 out of 6 preterm samples (66%) and misclassifying 3 of 18 full-term samples (17%) (Figure 1). Moreover, this independent validation cohort showed that it is possible to discriminate spontaneous preterm from full-term pregnancies up to 2 months in advance of labor with an AUC of 0.74 (Figure 3C). Several of the genes used to predict spontaneous preterm delivery were also individually significantly more highly expressed in women who delivered preterm (FDR ≤ 5%, Hedge’s g ≥ 0.8), demonstrating the robustness of their effect (Figure 3B). Our evidence suggests that the genes associated with spontaneous preterm birth are distinct from those found to be most predictive for GA and full-term delivery.

The cfRNA results can be compared attempts to estimate preterm risk using mass spectroscopic measurements of the ratio of two proteins (SHBG and IBP4) (*25*). Our method not only yields higher mean accuracy for comparable sample sizes in the validation cohorts (AUC=0.74 (cfRNA), AUC=0.67(IBP4/SHBG)), but also has several advantages, specifically broader applicability and cost. The mass spectroscopic approach is only applicable in a very narrow two week gestational interval and only on a limited range of body mass indices (BMI). Moreover, at a fraction of the cost of mass spectrometry, PCR can easily be applied to resource-challenged settings where its impact may be most direct.

In summary, we have described a molecular clock of pregnancy development that reflects a roadmap of placental and fetal gene expression, and enables prediction months in advance of time to delivery, GA, and expected delivery date with comparable accuracy to ultrasound. Because this model is derived from cfRNA measurements, it has several advantages, namely cost and applicability later during pregnancy. At a fraction of the cost of ultrasound, cfRNA measurements can be easily applied to resource-challenged settings. Even in countries that regularly use ultrasound, cfRNA presents an attractive, accurate alternative to ultrasound, especially during the second and third trimesters. Predicting expected delivery dates using cfRNA improves during gestation as opposed to ultrasound predictions, which deteriorate from 15 (T2) to 27 (T3) day estimates of delivery (*26*). We expect that this clock will also be useful for discovering and monitoring fetuses with congenital defects that can be treated *in utero*, which represents a rapidly growing area in maternal-fetal medicine. Furthermore, we investigated our model’s application to spontaneous preterm delivery, and discovered and validated a separate set of 10 biomarkers, suggesting that the physiology of preterm delivery is distinct from normal development, forming the basis for a screening test for assessing risk of spontaneous preterm birth.

## Acknowledgements

This work was supported by the Bill and Melinda Gates Foundation, March of Dimes Prematurity Research Center at Stanford University, March of Dimes Prematurity Initiative Grant at the University of Pennsylvania, and the Chan Zuckerberg Biohub. We thank Derek Croote and Mark Kowarsky for helpful discussions.

